# Preliminary genomic data on five *Orientia tsutsugamushi* strains isolated in Vellore, India

**DOI:** 10.1101/2025.03.07.642006

**Authors:** Janaki Kumaraswamy, Agilandeeswari Kirubanathan, Karthik Gunasekaran, KPP Abhilash, John Antony Jude Prakash

**Affiliations:** Department of Clinical Microbiology, Christian Medical College, Vellore, Tamil Nadu, India; Department of General Medicine, Christian Medical College, Vellore, Tamil Nadu, India; Emergency Medicine, Christian Medical College, Vellore, Tamil Nadu, India

**Keywords:** *Orientia tsutsugamushi*, whole genome sequencing, phylogeny, scrub typhus, India

## Abstract

**Introduction:** *Orientia tsutsugamushi*, the causative agent of scrub typhus, can be isolated in Vero or L929 cells and has a small genome (2–2.5 Mb). However, genome assembly is challenging due to the presence of host DNA contamination and a high proportion of repeat regions (up to 51%). Current global data includes 11 fully annotated genomes, with none from India. Here, we present the first whole-genome sequences of *O. tsutsugamushi* from India.

**Methods:** Five *O. tsutsugamushi* strains were cultured in Vero cells and confirmed by 47kDa real-time PCR. Genomic DNA was extracted after removal of host DNA and sequencing libraries were prepared. Whole-genome sequencing was performed using the PacBio Sequel II system in CCS/HiFi mode. The raw reads were assembled using Flye, and genome completeness was assessed with QUAST and BUSCO. Annotation was performed using the NCBI PGAP pipeline and comparative genome analysis by Roary. Phylogenetic analysis was based on the full-length 56kDa gene, which contains four variable domains.

**Results:** We report five complete genomes of *O. tsutsugamushi*, four of which are circular and one linear. Genome sizes range from 2.1 to 2.4 Mb. The total number of predicted genes falls between 2,379 and 2,715, with an average of 1,824 coding genes and 613 pseudogenes. Repeat regions constitute 53–59% of the genome, a higher proportion than previously reported. All five genomes have been submitted to NCBI GenBank (Accession Numbers: CP166954-58). Phylogenetic analysis based on the full-length 56kDa gene revealed that two strains belong to the Karp genogroup, two to Kato, and one to TA763.

**Conclusion:** This study presents the first whole-genome sequencing data of *O. tsutsugamushi* from India. Notably, the repetitive regions in these genomes are more extensive than previously reported. Further analyses with additional isolates are necessary to validate this observation.

Comprehensive phylogenomic studies, particularly to elucidate evolutionary dynamics and potential recombination events will provide further information.

## Introduction

*Orientia tsutsugamushi*, an obligate intracellular bacterium, is the etiological agent of scrub typhus, a vector-borne zoonotic disease (1). It is transmitted to humans, the dead-end host, through the bite of infected chiggers (2). Once inside the human host, *O. tsutsugamushi* invades macrophages and vascular endothelial cells, escapes from the phagosomes, and replicates within the cytosol. Clinically, scrub typhus presents with fever, eschar formation, rash and in severe cases, organ failure and death, if left untreated (3). The disease is effectively treated with anti-rickettsial agents such as doxycycline, chloramphenicol, and azithromycin; however, no licensed vaccine is currently available for prevention (4,5).

There is limited genomic data on *O. tsutsugamushi*, due to the challenges in isolation, purification, and sequencing (6). Amongst the 11 complete genomes (2.0–2.5 Mb) reported globally (https://www.ncbi.nlm.nih.gov/datasets/genome/?taxon=784&assembly_level=2:3), there is none from India (Prakash JAJ, personal communication). As an obligate intracellular parasite, *O. tsutsugamushi* cannot be cultured in conventional media and requires propagation in Vero or L929 cell lines (5,7,8). However, purifying *O. tsutsugamushi* from infected Vero cells is crucial for genome assembly, including gene annotation and comparative genomics (7).

The *O. tsutsugamushi* genome is highly repetitive, with repeat regions comprising 33–51% of the genome, including numerous pseudogenes, making whole-genome assembly particularly challenging (9–11). Long-read sequencing platforms such as PacBio and Oxford Nanopore Technologies have proven useful in resolving these repetitive regions, with very good capability for genome assembly reported for PacBio (12).

In this study, we report the whole-genome sequences (WGS) of five *Orientia tsutsugamushi* strains isolated in Vellore, Tamil Nadu, India using the PacBio Sequel II system.

## Methods

### Bacterial propagation

The five *Orientia tsutsugamushi* strains (JJOtsu1, JJOtsu5-8) isolated from scrub typhus patients were maintained in Vero cell lines (National Centre for Cell Science, Pune, Maharastra, India) using DMEM (Himedia, Thane, Maharashtra, India) supplemented with 2% FBS (Merck, Rahway, NJ, USA). The isolates were expanded in 75 cm^2^ flasks (Tarsons, Kolkata, West Bengal, India) and cultured for 10 days. Bacterial growth was confirmed by cytopathic effect observation and 47kDa real-time PCR (qPCR), as previously described (13).

### Host Genome Removal from *Orientia tsutsugamushi*-Infected Vero Cells

Bead beating and high-speed centrifugation (25,000g) methods were tried to purify *O. tsutsugamushi*, but the outcomes were unsatisfactory (Refer Supplementary Data). Therefore, an in-house approach (used for disrupting tissue samples sent for bacterial cultures) was adopted, and the results were satisfactory. The procedure is outlined below (in brief).

Infected cells were detached using a sterile scraper and lysed by vortexing with 5.3 mm sterile glass beads (The Science House, Chennai, India) for 1 minute. The lysate was centrifuged at 300g for 3 minutes at room temperature to pellet host cell debris. The supernatant was then filtered through a 5 µm syringe filter (Micro Separations, Ghaziabad, Uttar Pradesh, India) to remove host cell nuclei. Bacterial cells were pelleted by centrifugation at 14,000g for 10 minutes at room temperature. The pellet was resuspended in 200 µl of phosphate-buffered saline (PBS) and immediately processed for DNA extraction.

### DNA Extraction

DNA was extracted from the reconstituted pellet (200 µl) using DNeasy Blood and Tissue kit (Qiagen, Hilden, Germany) as per the manufacturer’s instructions. The quantity and purity of extracted DNA was measured using Qubit 4.0 fluorometer and Nano-Drop 2000 (Thermo Fisher Scientific, Waltham, MA, USA). DNA integrity and size were estimated using 1% agarose gel and by Agilent FEMTO pulse analyser (Agilent Technologies, Santa Clara, CA, USA). The proportion of host (Vero cell) to bacterial DNA was measured using Real-time PCR targeting beta actin gene (Vero) and 47kDa (*Orientia tsutsugamushi*) gene respectively (13,14).

### Library construction and Sequencing

DNA shearing was performed on Megaruptor 3 (Diagenode, Seraing, Belgium) followed by end repair/A-tailing. The library was constructed using the SMRTbell Express template Preparation Kit 3.0 (Pacific Biosciences, Menlo Park, CA, USA) as per manufacturers’ protocol. The prepared library was purified using AMPure PB beads (Pacific Biosciences, Menlo Park, CA, USA). All the bacterial libraries were pooled according to volumes provided by Microbial Multiplexing Calculator (Pacific Biosciences, Menlo Park, CA, USA). The size distribution of final pooled SMRTbell library was determined by Agilent FEMTO pulse analyser (Agilent Technologies, Santa Clara, CA, USA). The size selected SMRT libraries were purified and subjected to primer annealing and polymerase binding using Sequel II binding kit 3.2 (Pacific Biosciences, Menlo Park, CA, USA) to prepare bound complex. About 90 pM of this library was loaded on to SMRTcell containing 8M ZMW and sequenced in PacBio Sequel II system (Pacific Biosciences, Menlo Park, CA, USA) in CCS/HiFi mode.

### Bioinformatic analysis

Raw sub-reads generated were converted to HiFi reads using circular consensus sequencing (Version-6.2.0) and assembled using Flye (Version - 2.9.3-b1797), a denovo genome assembler (15). The quality and completeness of the assembly was evaluated by QUAST (QUality ASsessment Tool for genome assemblies) (16) and BUSCO (Benchmarking Universal Single-Copy Orthologs), a quantitative assessment tool for genome assemblies based on evolutionarily informed expectations of gene content from near-universal single-copy orthologs (17). The circular genomes were verified by Bandage: Interactive visualisation of denovo assembly (18). The NCBI PGAP pipeline Version 6.8 (19) was used to annotate the genomes. The repeat region in the assembled genomes were identified using Repeat modeler Version 2.0.5 (20).

### Phylogenetic analysis

A phylogenetic tree was constructed using complete 56kDa gene. Alignment was performed using Clustal Omega(21) with 47 reference sequences (including complete 56kDa gene from 11 reference genome) retrieved from GenBank-NCBI. The phylogenetic tree was established for complete 56kDa gene using IQTREE software: A fast and effective stochastic algorithm for estimating maximum likelihood phylogenies (22).

### Results

The method used for purification of *Orientia tsutsugamushi* showed efficient depletion of Vero cell (host cell) DNA. A Ct value difference of 15 (5-log) was observed between bacteria to host, suggesting >99% depletion of host (Vero) genome refer **Table 1**.

**Table 1.** Ct value of host (Vero) specific β actin gene and *Orientia tsutsugamushi* (O.t) specific 47kDa gene.

| S no. | Strain | Ot specific 47kDa (Ct value) | Vero specific $\beta$ actin gene (Ct value) |
| --- | --- | --- | --- |
| 1 | JJOtsu1 | 14 | 30 |
| 2 | JJOtsu5 | 15 | 28 |
| 3 | JJOtsu6 | 16 | 29 |
| 4 | JJOtsu7 | 18 | 29 |
| 5 | JJOtsu8 | 14 | 31 |

PacBio generated five single contig assembly with four being circular and one linear genome. The genomes have been submitted to NCBI Genbank with accession number CP166954-58. The genome size ranges from 2.1 to 2.4 Mb. The number of predicted genes ranges from 2379 to 2715 and the repeat regions range from 52.88 to 58.89 %. A total of 657 genes were present in all the five genomes and constitute the core genome amongst our isolates. The complete annotation summary is as given in **Table 2**. Phylogenetic analysis of complete 56kDa gene sequence showed that JJOtsu1 and JJOtsu6 clustered into Kato A genotype (Kato genogroup), JJOtsu5 and JJOtsu7 clustered into Karp A genotype (Karp genogroup) and JJOtsu8 belongs to TA763 B genotype (TA763 genogroup) refer **Figure 1**.

**Table 2.** *Orientia tsutsugamushi* WGS summary.

| Strain name | JJOtsu1 | JJOtsu5 | JJOtsu6 | JJOtsu7 | JJOtsu8 |
| --- | --- | --- | --- | --- | --- |
| Accession number | CP166954 | CP166955 | CP166956 | CP166957 | CP166958 |
| Genogroup | Kato | Karp | Kato | Karp | TA763 |
| Length of 56kDa gene | 1569 | 1608 | 1569 | 1605 | 1605 |
| Number of contigs | 1 | 1 | 1 | 1 | 1 |
| Total length | 2179789 | 2446845 | 2284757 | 2183885 | 2344138 |
| Assembly level | Complete | Complete | Complete | Complete | Complete |
| Assembly Type | Circular | Circular | Circular | Linear | Circular |
| N50 | 2179789 | 2446845 | 2284757 | 2183885 | 2344138 |
| N90 | 2179789 | 2446845 | 2284757 | 2183885 | 2344138 |
| Genome coverage (X) | 1168 | 1636 | 1343 | 1778 | 1408 |
| GC% | 30.53 | 30.71 | 30.63 | 30.48 | 30.57 |
| Repeat region (%) | 54.71 | 58.89 | 56.90 | 52.88 | 56.69 |
| Genes (Total) | 2381 | 2715 | 2468 | 2379 | 2473 |
| CDS (Total) | 2341 | 2675 | 2429 | 2339 | 2432 |
| Gene (coding) | 1764 | 1899 | 1848 | 1767 | 1850 |
| rRNA (5s, 16s, 23s) | 3 | 3 | 3 | 3 | 3 |
| tRNA | 34 | 34 | 33 | 34 | 35 |
| ncRNA | 3 | 3 | 3 | 3 | 3 |
| Pseudogene | 577 | 776 | 581 | 572 | 582 |

**Legend:** rRNA: ribosomal RNA, tRNA: transfer RNA, ncRNA: non-coding RNA, GC: Guanine-Cytosine, CDS: Coding sequence.

**Figure 1:**
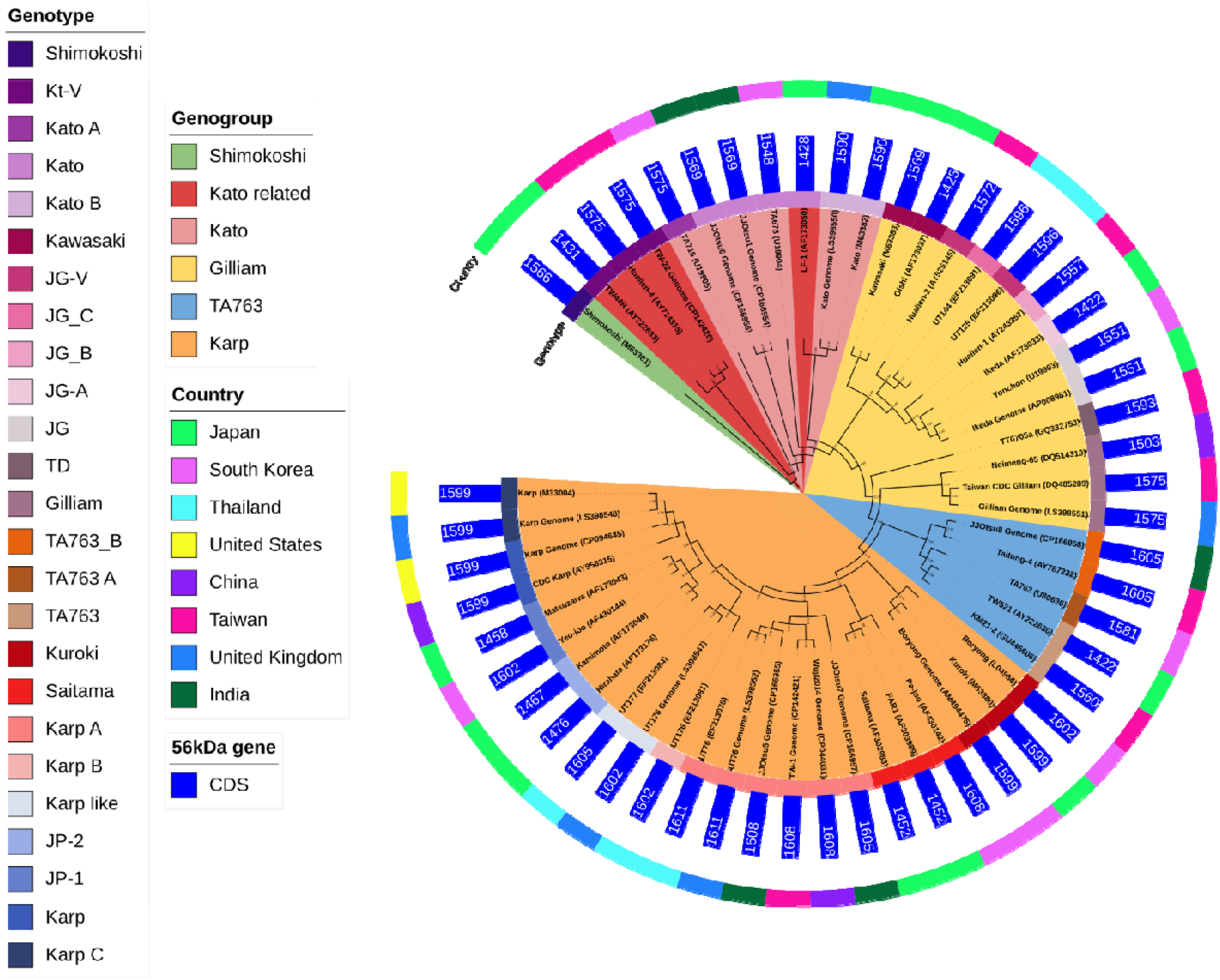
Phylogenetic tree of *Orientia tsutsugamushi* strains based on complete 56kDa gene sequence.

## Discussion

### DNA Quality and Host Interference

One of the critical challenges in sequencing *Orientia tsutsugamushi* is the presence of a significant amount of host DNA, as this bacterium propagates inside host cells (in this case, **Vero cells**). The Vero genome is much larger than that of the bacterium (3 GB vs. 2–2.5 Mb) (23), which leads to a high proportion of host DNA contaminating the sample and interfering with the sequencing process. This study employed a purification protocol that achieved >99% depletion of the host genome. By minimizing host contamination, this technique allows for enrichment of the bacterial genome, crucial for obtaining high-quality sequencing reads necessary for assembly.

This method stands out because it does not require expensive equipment, making it more accessible compared to other high-cost alternatives. Although it is labour-intensive, it strikes a balance between cost-effectiveness and efficiency.

### Challenges in Assembling the *O. tsutsugamushi* Genome

The genome of *O. tsutsugamushi* is highly repetitive, with repetitive regions making up as much as 51% of the genome. Repetitive DNA sequences present a significant challenge for genome assembly (11), as they can cause difficulties in accurately aligning sequencing reads and producing complete, contiguous sequences. Historically, assembling the genome of *O. tsutsugamushi* has been difficult, and older methods like **Bacterial Artificial Chromosome (BAC) cloning** and **Sanger sequencing** were used (9,24). While these methods provided valuable data, they were time-consuming and limited in resolving repetitive regions.

The PacBio Sequel II system, with its ability to produce high accuracy long reads is particularly beneficial for sequencing repetitive regions and assembling complex genomes like that of *O. tsutsugamushi*. Batty et al., 2018 used PacBio RS II sequencing which were polished using Illumina sequencing to obtain six complete genome of *O. tsutsugamushi*.(11). We used the latest generation PacBio sequel II sequencer to generate long sequences of high accuracy, which resulted in obtaining single contig genomes for our five *O. tsutsugamushi strains* (Vellore isolates) by de-novo assembly. The core genome analysis of these Vellore isolates showed 657 genes, and is consistent with that reported by Batty *et al*., in 2018. The repeat regions in our five strains ranged from 52.9% to 58.9%, this is higher than the 33% to 51% reported in the literature (11)

### Global Data and Genomic Sequences

At the time of the study, there were only **11 well-annotated whole genome sequences** of *O. tsutsugamushi* available globally, which limits the scope for comprehensive **WGS-based phylogenetic analysis**. The addition of the five Vellore isolates genomes in this study increases the available genomic data, but more strains need to be sequenced to make robust phylogenetic comparisons and understand the evolutionary relationships between strains.

The **TSA 56 gene** (also known as the **56 kDa antigen gene**) is traditionally used as a marker for identifying the **genogroup** and **genotype** of *O. tsutsugamushi*. Variations in its four variable domains (VD I – VD IV) help differentiate between genotypes. In this study, **47 complete 56kDa gene sequences** were used to generate a phylogenetic tree, which helped clarify the relationships between different *O. tsutsugamushi* strains. Amongst the five isolates analysed, two belonged to Kato A genotype (Kato genogroup), two belonged to Karp A genotype (Karp genogroup) and one belongs to TA763 B genotype (TA763 genogroup). This phylogenetic analysis using the TSA 56 gene provides insight into the genetic diversity and the distribution of different genotypes within *O. tsutsugamushi*. It also offers a foundation for understanding the evolutionary relationships and potential geographic variation within the species, which is important for epidemiological studies.

## Conclusion

We report for the first time from India five well annotated and complete genomes of *Orientia tsutsugamushi* strains (includes two Kato, two Karp and one TA763 genogroups). This study represents an important step forward in understanding the genomics of *O. tsutsugamushi*, with the use of the PacBio Sequel II system providing a more detailed and accurate genomic map. Further, phylogenomic analysis is necessary to determine evolutionary relationships, recombination events, mobilome and synteny of Indian strains versus those found in other regions.

## Acknowledgements

We acknowledge Nucleome Informatics Private, Limited, Hyderabad, Telangana, India for PacBio sequencing and help in data curation.

## Data availability

This whole-genome sequencing project has been archived in the NCBI database under the accession number <u>PRJNA1141359</u>.

## Ethics approval

The study was approved by the Institutional Review Board (IRB) & Ethics Committee vide IRB Min No. 11942 dated 27th March 2019; National Reference Centre for Rickettsial Diseases IRB Min. no. 0624131 dated 12.06.2024

## Author contribution statement

Study design: J.K., A.K., K.G., K.P.P., J.A.J.P.; Data collection: J.K., A.K., K.G.; Data analysis and interpretation: J.K., A.K., K.G., K.P.P., J.A.J.P.; Drafting the manuscript: J.K., A.K.; Critical revision of the manuscript: K.G., K.P.P., J.A.J.P.; Supervision and funding acquisition: J.A.J.P.

## References

1. Watt G, Parola P. Scrub typhus and tropical rickettsioses. Curr Opin Infect Dis. 2003 Oct;16(5):429–36.

2. Demma LJ, McQuiston JH, Nicholson WL, Murphy SM, Marumoto P, Sengebau-Kingzio JM, et al. Scrub Typhus, Republic of Palau. Emerg Infect Dis. 2006 Feb;12(2):290–5.

3. Ko Y, Choi JH, Ha NY, Kim IS, Cho NH, Choi MS. Active Escape of Orientia tsutsugamushi from Cellular Autophagy. Infect Immun. 2013 Feb;81(2):552–9.

4. Walker DH. Scrub Typhus — Scientific Neglect, Ever-Widening Impact. N Engl J Med. 2016 Sep 8;375(10):913–5.

5. Luce-Fedrow A, Lehman ML, Kelly DJ, Mullins K, Maina AN, Stewart RL, et al. A Review of Scrub Typhus (Orientia tsutsugamushi and Related Organisms): Then, Now, and Tomorrow. Trop Med Infect Dis. 2018 Jan 17;3(1):8.

6. Elliott I, Thangnimitchok N, de Cesare M, Linsuwanon P, Paris DH, Day NPJ, et al. Targeted capture and sequencing of Orientia tsutsugamushi genomes from chiggers and humans. Infect Genet Evol. 2021 Jul 1;91:104818.

7. Giengkam S, Blakes A, Utsahajit P, Chaemchuen S, Atwal S, Blacksell SD, et al. Improved Quantification, Propagation, Purification and Storage of the Obligate Intracellular Human Pathogen Orientia tsutsugamushi. PLoS Negl Trop Dis. 2015 Aug 28;9(8):e0004009.

8. Ming DK, Phommadeechack V, Panyanivong P, Sengdatka D, Phuklia W, Chansamouth V, et al. The Isolation of Orientia tsutsugamushi and Rickettsia typhi from Human Blood through Mammalian Cell Culture: a Descriptive Series of 3,227 Samples and Outcomes in the Lao People’s Democratic Republic. J Clin Microbiol. 2020 Nov 18;58(12):e01553–20.

9. Nakayama K, Yamashita A, Kurokawa K, Morimoto T, Ogawa M, Fukuhara M, et al. The Whole-genome Sequencing of the Obligate Intracellular Bacterium Orientia tsutsugamushi Revealed Massive Gene Amplification During Reductive Genome Evolution. DNA Res. 2008 Aug 1;15(4):185–99.

10. Cho NH, Kim HR, Lee JH, Kim SY, Kim J, Cha S, et al. The Orientia tsutsugamushi genome reveals massive proliferation of conjugative type IV secretion system and host– cell interaction genes. Proc Natl Acad Sci. 2007 May 8;104(19):7981–6.

11. Batty EM, Chaemchuen S, Blacksell S, Richards AL, Paris D, Bowden R, et al. Long-read whole genome sequencing and comparative analysis of six strains of the human pathogen Orientia tsutsugamushi. PLoS Negl Trop Dis. 2018 Jun 6;12(6):e0006566.

12. Oehler JB, Wright H, Stark Z, Mallett AJ, Schmitz U. The application of long-read sequencing in clinical settings. Hum Genomics. 2023 Aug 8;17(1):73.

13. Kim DM, Park G, Kim HS, Lee JY, Neupane GP, Graves S, et al. Comparison of Conventional, Nested, and Real-Time Quantitative PCR for Diagnosis of Scrub Typhus. J Clin Microbiol. 2011 Feb;49(2):607–12.

14. Design and development of a simple method for the detection and quantification of residual host cell DNA in recombinant rotavirus vaccine. Mol Cell Probes. 2021 Feb 1;55:101674.

15. Kolmogorov M, Yuan J, Lin Y, Pevzner PA. Assembly of long, error-prone reads using repeat graphs. Nat Biotechnol. 2019 May;37(5):540–6.

16. Gurevich A, Saveliev V, Vyahhi N, Tesler G. QUAST: quality assessment tool for genome assemblies. Bioinformatics. 2013 Apr 15;29(8):1072–5.

17. Manni M, Berkeley MR, Seppey M, Zdobnov EM. BUSCO: Assessing Genomic Data Quality and Beyond. Curr Protoc. 2021;1(12):e323.

18. Wick RR, Schultz MB, Zobel J, Holt KE. Bandage: interactive visualization of de novo genome assemblies.

19. Tatusova T, DiCuccio M, Badretdin A, Chetvernin V, Nawrocki EP, Zaslavsky L, et al. NCBI prokaryotic genome annotation pipeline. Nucleic Acids Res. 2016 Aug 19;44(14):6614–24.

20. Flynn JM, Hubley R, Goubert C, Rosen J, Clark AG, Feschotte C, et al. RepeatModeler2 for automated genomic discovery of transposable element families. Proc Natl Acad Sci U S A. 2020 Apr 28;117(17):9451–7.

21. Sievers F, Wilm A, Dineen D, Gibson TJ, Karplus K, Li W, et al. Fast, scalable generation of high-quality protein multiple sequence alignments using Clustal Omega. Mol Syst Biol. 2011 Oct 11;7:539.

22. Nguyen LT, Schmidt HA, von Haeseler A, Minh BQ. IQ-TREE: a fast and effective stochastic algorithm for estimating maximum-likelihood phylogenies. Mol Biol Evol. 2015 Jan;32(1):268–74.

23. Sène MA, Kiesslich S, Djambazian H, Ragoussis J, Xia Y, Kamen AA. Haplotype-resolved de novo assembly of the Vero cell line genome. NPJ Vaccines. 2021 Aug 20;6:106.

24. Giengkam S, Kullapanich C, Wongsantichon J, Adcox HE, Gillespie JJ, Salje J. Orientia tsutsugamushi: comprehensive analysis of the mobilome of a highly fragmented and repetitive genome reveals the capacity for ongoing lateral gene transfer in an obligate intracellular bacterium. mSphere. 2023 Oct 18;8(6):e00268–23.

25. Long J, Wei Y, Tao X, He P, Xu J, Wu X, Zhu W, Chen K, Yang Z. Representative Genotyping, Recombination and Evolutionary Dynamics Analysis of TSA56 Gene Segment of Orientia tsutsugamushi. Front Cell Infect Microbiol. 2020 Aug 5;10:383.

26. Kumaraswamy J, Govindasamy P, Nagarajan LS, Gunasekaran K, Abhilash KPP, Prakash JAJ. Genotyping of Orientia tsutsugamushi circulating in and around Vellore (South India) using TSA 56 gene. Indian J Med Microbiol. 2024;47:100483.

