## Supplementary data for "Preliminary genomic data on five *Orientia tsutsugamushi* strains isolated in Vellore, India"

**Host lysis**

1. **Bead beating and DNase**

The infected cells were scrapped and lysed using bead beating lysis system (Fastprep-24, MP Biomedicals, Santa Ana, CA, USA) with beads (Lysing matrix D, MP Biomedicals, Santa Ana, CA, USA) operated at power 6 for 40 seconds. The lysate was centrifuged at 300g for 3minutes to pellet host cell debris. The supernatant was passed through 2-micron filter followed by addition RQ1 RNase-Free DNase (Promega corporation, Madison, WI, USA) to remove host cell DNA. The bacterial cells were pelleted by centrifugation at 14000g for 10 minutes. The pellet was resuspended with 1x PBS (Phosphate Buffered Saline) followed by DNA extraction using DNeasy Blood and Tissue kit (Qiagen, Hilden, Germany) as per the manufacturer’s instructions.

2. **High speed centrifugation**

The infected cells were scraped and resuspended in 4ml TS buffer (33mM Tris, 250mM sucrose, pH 7.4) and lysed by passing through 22-gauge syringe (10 times). The lysate was centrifuged at 300g for 10 mins. The supernatant was added on to 30% Percoll solution (Sigma -Aldrich, ) and centrifuged at 25000g for 60 minutes. The bottom layer containing the bacteria was removed and subjected to DNA extraction using DNeasy Blood and Tissue kit (Qiagen, Hilden, Germany) as per the manufacturer’s instructions.

**Real-time PCR**

The extracted DNA was quantified using Nano-Drop 2000 (Thermo Fisher Scientific, Waltham, MA, USA) and subjected to real-time PCR targeting beta actin gene (Vero) and 47kDa (*Orientia tsutsugamushi*) gene respectively.

**Result**

The difference between host and bacteria was 10 Ct in the bead beating method and 14-18 Ct in the high-speed centrifugation method. The DNA concentration in both procedures was relatively low, which is insufficient for sequencing (Refer table 1 below for details). The minimal amount needed for obtaining best sequence results using PacBio sequel II is 1000ng.

**Table 1. Ct value and DNA concentration**

| **Method** | **Strain** | **Ct Value** | | **DNA concentration**  **(ng/µl)** | **Total DNA (100µl)*** |
| --- | --- | --- | --- | --- | --- |
|  |  | **O. t specific 47kDa gene** | **Vero specific β actin gene** |  |  |
| Bead beating and DNase | JJOtsu1 | 20.3 | 30.1 | 1.4 | 140 |
|  | JJOtsu5 | 17.1 | 27.4 | 7.9 | 790 |
| High speed centrifugation | JJOtsu1 | 18 | 32 | 3.9 | 390 |
|  | JJOtsu5 | 20 | 38 | 0.9 | 90 |

*100µl elution volume
